# A Potent and Selective CDKL5/GSK3 Chemical Probe is Neuroprotective

**DOI:** 10.1101/2023.02.09.527935

**Authors:** Han Wee Ong, Yi Liang, William Richardson, Emily R. Lowry, Carrow I. Wells, Xiangrong Chen, Margaux Silvestre, Kelvin Dempster, Josie A. Silvaroli, Jeffery L. Smith, Hynek Wichterle, Navjot S. Pabla, Sila K. Ultanir, Alex N. Bullock, David H. Drewry, Alison D. Axtman

**Affiliations:** Structural Genomics Consortium, UNC Eshelman School of Pharmacy, University of North Carolina at Chapel Hill, Chapel Hill, North Carolina, 27599, United States of America; Centre for Medicines Discovery, Nuffield Department of Medicine, University of Oxford, Oxford, OX3 7DQ, United Kingdom; Department of Pathology and Cell Biology, Columbia University Irving Medical Center, New York, New York, 10032, United States of America; The Project ALS Therapeutics Core, Columbia University Irving Medical Center, New York, New York, 10032, United States of America; Kinases and Brain Development Laboratory, The Francis Crick Institute, London, NW1 1AT, United Kingdom; Division of Pharmaceutics and Pharmacology, College of Pharmacy and Comprehensive Cancer Center, The Ohio State University, Columbus, Ohio, 43210, United States of America; Departments of Pathology and Cell Biology, Neurology, Neuroscience, Rehabilitation and Regenerative Medicine, Columbia University Irving Medical Center, New York, New York, 10032, United States of America; Center for Motor Neuron Biology and Disease, Columbia University Irving Medical Center, New York, New York, 10032, United States of America; Columbia Stem Cell Initiative, Columbia University Irving Medical Center, New York, New York, 10032, United States of America; UNC Lineberger Comprehensive Cancer Center, School of Medicine, University of North Carolina at Chapel Hill, Chapel Hill, North Carolina, 27599, United States of America

**Keywords:** CDKL5, GSK3α, GSK3β, chemical probe, neuroprotective, kinase, crystal structure

## Abstract

Despite mediating several essential processes in the brain, including during development, cyclin-dependent kinase-like 5 (CDKL5) remains a poorly characterized human protein kinase. Accordingly, its substrates, functions, and regulatory mechanisms have not been fully described. We realized that availability of a potent and selective small molecule probe targeting CDKL5 could enable illumination of its roles in normal development as well as in diseases where it has become aberrant due to mutation. We prepared analogs of AT-7519, a known inhibitor of several cyclin dependent and cyclin-dependent kinase-like kinases that has been advanced into Phase II clinical trials. We identified analog **2** as a highly potent and cell-active chemical probe for CDKL5/GSK3 (glycogen synthase kinase 3). Evaluation of its kinome-wide selectivity confirmed that analog **2** demonstrates excellent selectivity and only retains GSK3α/β affinity. As confirmation that our chemical probe is a high-quality tool to use in directed biological studies, we demonstrated inhibition of downstream CDKL5 and GSK3α/β signaling and solved a co-crystal structure of analog **2** bound to CDKL5. A structurally similar analog (**4**) proved to lack CDKL5 affinity and maintain potent and selective inhibition of GSK3α/β. Finally, we used our chemical probe pair (**2** and **4**) to demonstrate that inhibition of CDKL5 and/or GSK3α/β promotes the survival of human motor neurons exposed to endoplasmic reticulum (ER) stress. We have demonstrated a neuroprotective phenotype elicited by our chemical probe pair and exemplified the utility of our compounds to characterize the role of CDKL5/GSK3 in neurons and beyond.

## INTRODUCTION

Cyclin-dependent kinase-like 5 (CDKL5) is an understudied, human serine/threonine kinase from the CMGC group of kinases. It is a member of the human CDKL family of kinases, which also includes CDKL1, CDKL2, CDKL3, and CDKL4.^1^ CDKL5 was included on the list of ‘dark kinases’ defined by the NIH for the Illuminating the Druggable Genome (IDG) program.^2^ CDKL5 is also known as serine/threonine kinase 9 (STK9).^3^ CDKL5 protein is the product of the X-linked gene of the same name.^4^

While CDKL5 is widely expressed in most tissues and cells, it is predominantly found in the brain.^5, 6^ This kinase is expressed abundantly in neurons within the brain, specifically in the cortex, hippocampus, striatum, and to a lesser extent in the cerebellum.^7-10^ It can localize to the nucleus, cytoplasm, cilia, centrosome, midbody, excitatory synapses, and/or dendritic branches, depending on its function and stage of development.^1, 3, 10-12^ In addition to a role in negatively influencing its catalytic activity, the C-terminal tail of CDKL5 has been implicated in localizing the protein to the cytoplasm when necessary.^10, 13^

CDKL5 is important for neuronal survival, proliferation, maturation, differentiation, and morphogenesis.^3, 4, 14^ It also contributes to dendritic morphogenesis and spine structure, synapse formation and activity, axon outgrowth, and cilia length.^1, 4, 5, 8, 12^ Many of these processes are impacted in neurodegenerative diseases, which are also characterized by endoplasmic reticulum (ER) stress.^15-17^ During development, CDKL5 aids in the establishment of the GABAergic network in the cerebellar cortex.^18^ It has also been implicated in regulating the behavior and morphology of nuclear speckles, which indirectly impacts pre-mRNA processing, and in cytokinesis.^5, 11^

Most literature for CDKL5 has focused on its important role in a rare and severe neurodevelopmental condition called CDKL5-deficiency disorder (CDD).^19, 20^ Patients with CDKL5 mutations were initially diagnosed with atypical Rett syndrome, autism, or West-syndrome due to the clinical similarities.^3, 11, 12, 21^ More than 70 different point mutations have been described, including missense mutations within the catalytic domain, nonsense mutations in the entire protein that cause its premature termination, frameshift mutations, and splice variants, and efforts have been made to correlate mutations with clinical outcomes.^21, 22^ Without active, full-length CDKL5, brain development and function are impaired.^8, 10, 22^ Accordingly, symptoms of CDD include early onset epileptic seizures before the age of 3 months, low muscle tone and orthopedic complications, impaired growth, developmental delays, intellectual disability, vision problems, sleep disturbances, difficulties associated with feeding and/or swallowing, gastrointestinal issues, and subtle facial, limb, and hand phenotype.^19, 23^ Although it is similar to Rett syndrome, it is classified as an independent condition rather than another variant.^23^ There is currently no cure and there are no specific treatments for CDD.

Despite its important role in brain function, the function of CDKL5 during brain development, its substrates, and the molecular mechanisms involved in its regulation remain largely uncharacterized.^14, 24^ Some substrates of CDKL5 include itself (auto-phosphorylation of Thr-Glu-Tyr motif), HDAC4, NGL-1, MAP1S, EB2, ARHGEF2, CEP131, DLG5, Dnmt1, ELOA, Sox9, and Amph1.^1, 6, 13, 24-26, 27, 28^ While there is varying degrees of validation of these proteins as physiological substrates of CDKL5, EB2 has been validated as a substrate by multiple groups and has been shown to be altered in CDD patient iPSC-derived neurons.^25, 29^ CDKL5 interacts with several proteins and pathways, including MeCP2, Dnmt1, Rac1, HIPK2/H2B, and PDK1/AKT/GSK3β. ^3-5, 9-11, 27, 30, 31^

There is crosstalk between CDKL5 and GSK3α/β. Loss of CDKL5 in a *CDKL5* knockout mouse model results in altered AKT/GSK3β signaling, resulting in deficient neuronal maturation and increased neuronal apoptosis, which have been attributed to increased activity of GSK3β.^4, 30^ *CDKL5* knockout mice also had impaired hippocampus-dependent memory.^4^ Evidence suggests that these alterations may underlie the developmental defects caused by loss of CDKL5.^30^ Importantly, this activation of GSK3β and its detrimental effects can be reversed via treatment with GSK3β inhibitors *in vivo*.^19, 20^ Accordingly, selective inhibition of GSK3 has proven neuroprotective *in vivo* and has been examined as a treatment option for neurodegenerative and psychiatric disorders.^32^

The co-crystal structure of the CDKL5 kinase domain bound to ASC67 was solved (PDB code: 4BGQ).^12^ ASC67 is closely related to a very promiscuous kinase inhibitor.^12, 33^ It is not specific to CDKL5. Although the authors of the crystallography paper note that the structure and inhibitor screens that they carried out suggest potential for generating isoform-selective inhibitors of the CDKL family, a potent and selective CDKL5 chemical probe has not been described.^12^

## RESULTS AND DISCUSSION

Aiming to better characterize the roles of CDKL5 in primary neurons, CDKL5/GSK3 chemical probe **2** was designed and synthesized to be used in cell-based studies. The suitability of **2** as a chemical probe for CDKL5 and GSK3 was investigated in binding assays, broad kinome screening, enzyme assays, co-crystallographic studies, and cell-based studies, including Western blot and an assay aimed at assessing its neuroprotective properties.

### Chemistry

We modified AT-7519, a compound that was developed via fragment-based X-ray crystallography and structure based drug design by Astex in 2008 as a cyclin dependent kinase 2 (CDK2) inhibitor.^34^ Although it was later found to inhibit many CDK and CDKL family members,^35^ AT-7519 has been advanced into multiple Phase I and Phase II clinical trials for advanced solid tumors, chronic lymphocytic leukemia (CLL), and mantle cell lymphoma, indicative of favorable properties.

To gain insight into those moieties on AT-7519 that are likely essential for binding to CDKL5, we docked AT-7519 to CDKL5. As shown in Figure 1, the pyrazole common to both ASC67 and AT-7519 makes key hydrogen bonding contacts with the hinge residues of CDKL5 (Glu90, Tyr91, and Val92 for ASC67). The 3-position amide was also proposed to hydrogen bond via its NH with the hinge residue Val92 of CDKL5 when AT-7519 was docked. An ionic interaction was also observed between the 3-position side chain piperidine nitrogen of AT-7519 with Glu98 of CDKL5.^12^ An interaction between a chloro-group on the 2,6-dichlorobenzene ring of AT-7519 with Lys42 of CDKL5 was noted, which mimics the hydrogen bond formed between Lys42 and the nitrile group of ASC67. This docking exercise suggested that to make CDKL5 active compounds we should not modify the pyrazole core or 3-position amide, should keep a heteroatom in the 3-position side chain, and should maintain halogenation of the 4-position side chain. It also suggested that the pocket would likely not accommodate chain extension at the 3-position without steric clash with residues in the ATP pocket.

**Figure 1.**
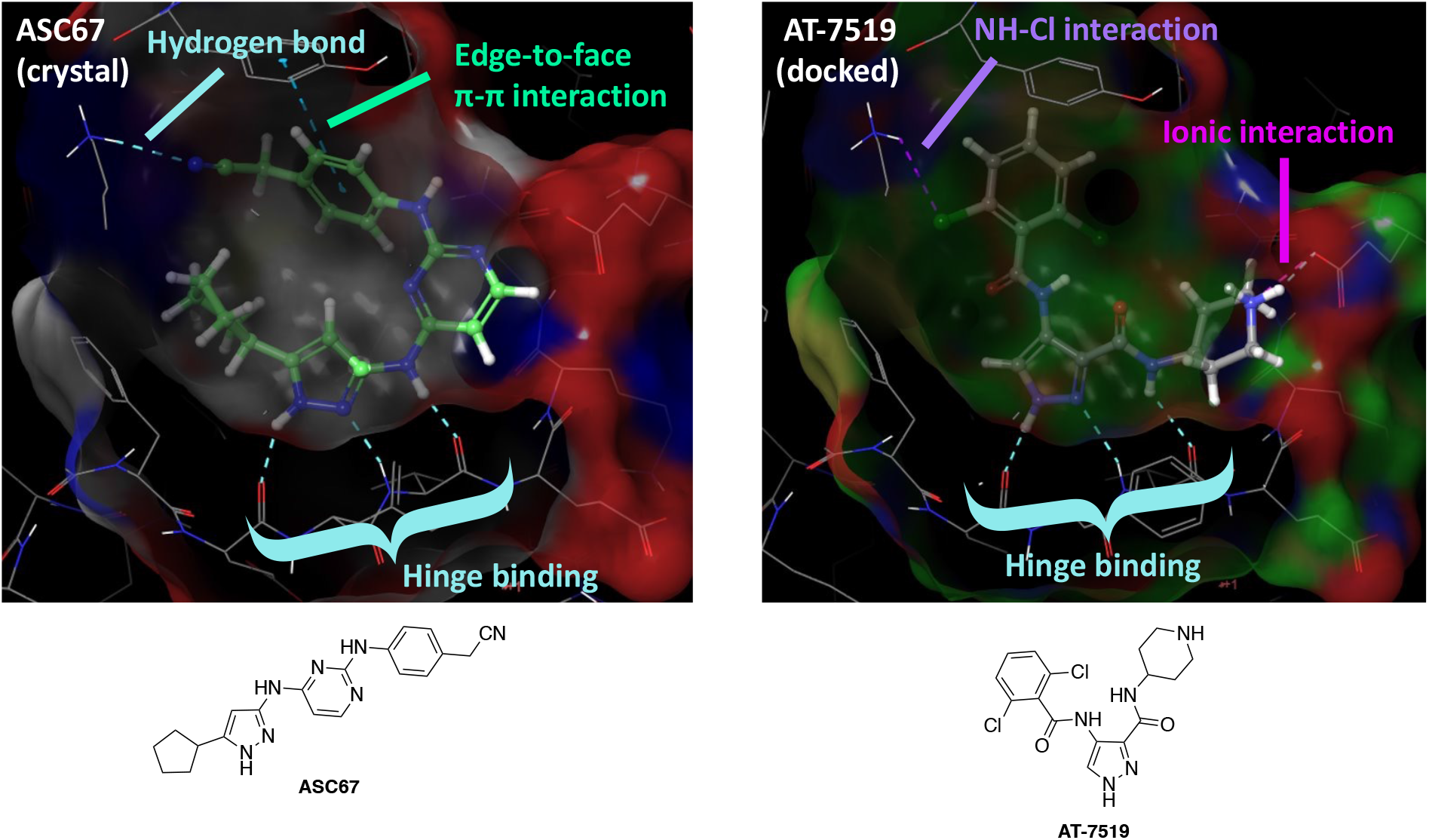
Comparison of CDKL5 co-crystal structure with ASC67 (left, PDB code: 4BGQ) and docking of AT-5419 (right). Key interactions, including the critical hinge binding interactions, are highlighted.

**Scheme 1.**
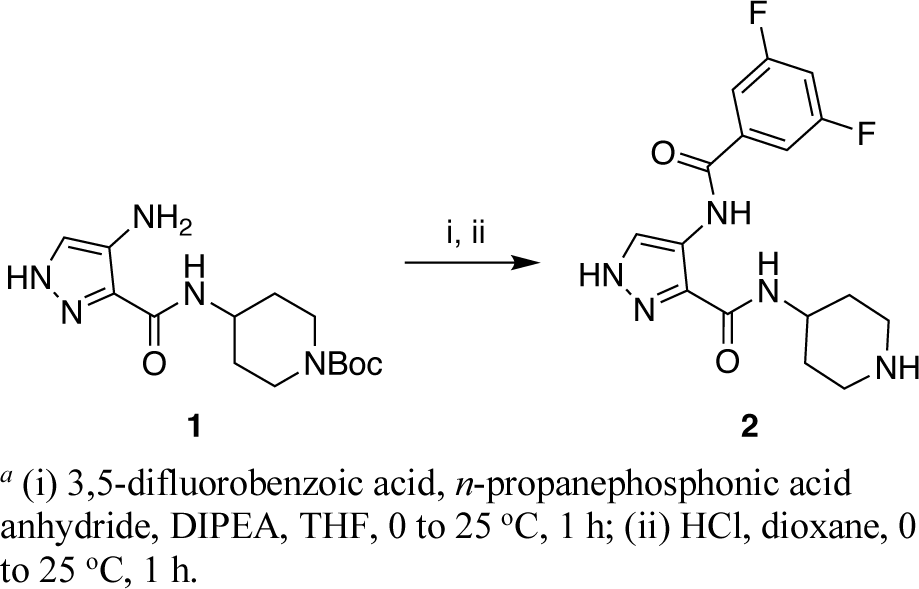
Synthesis of chemical probe compound (**2**)^*a*^

**Scheme 2.**
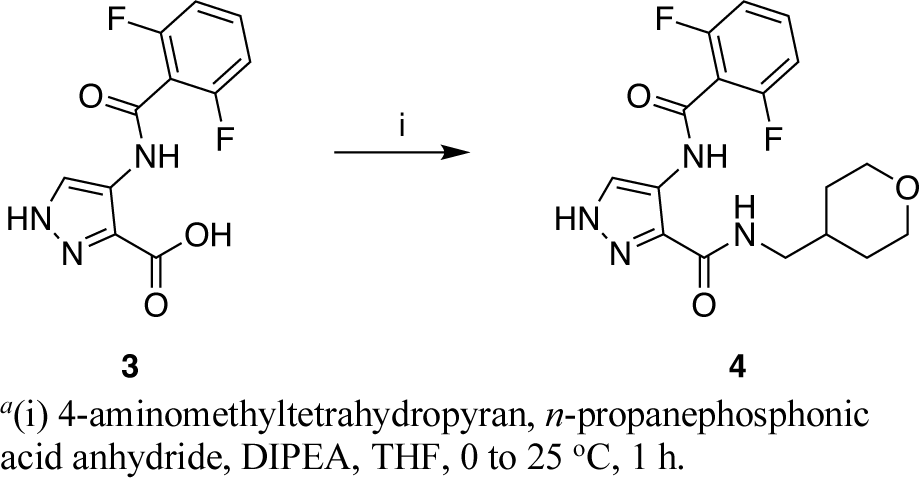
Synthesis of negative control compound (**4**)^*a*^

Consideration of the more than 130 unpublished AT-7519 analogs that we have made to date as well as published compounds have informed structure–activity relationships (SAR) for CDKL5. Figure 2 summarizes some important SAR determined through analog design. At the 4-position of the pyrazole core, mono- and di-fluorinated or chlorinated benzene rings proved the most efficacious for CDKL5 and demonstrated similar affinity to AT-7519. These halogenated rings resulted in better affinity to CDKL5 when compared to analogs with a cyanophenyl, pyridine, pyrimidine, or methylpyrazole at that same position. Alterations to the 3-position piperidine, including with 4-methylpiperidine, 3-pyrrolidine, or uncharged hydrophobic groups, all resulted in loss of CDKL5 affinity, suggesting that it was the best choice.

**Figure 2.**
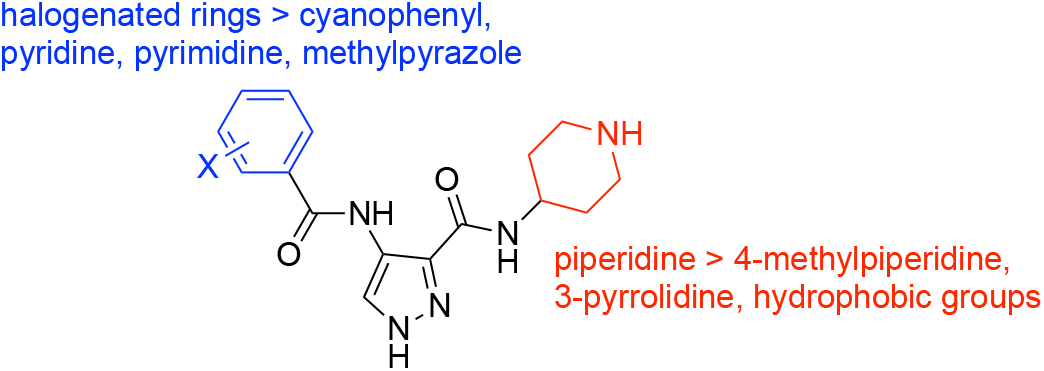
SAR summary for AT-7519 analogs.

Based on these SAR findings, we designed compound **2**. This analog couples the heightened CDKL5 affinity of a di-fluorinated benzene ring on the 4-position of the pyrazole core with the robust CDKL5 affinity of a 3-position piperidine on the pyrazole core. We hypothesized that these modifications would be synergistic and result in a potent inhibitor of CDKL5. The route summarized in Scheme 1, which involves amide bond formation followed by acid-mediated deprotection of the piperidine, was used to synthesize compound **2**. For a negative control compound, we designed analog **4** with a methylene inserted between the 3-position amide and heteroatom-containing ring on the pyrazole core. We hypothesized based on Figure 1 that the ATP binding pocket would not accommodate the longer chain since it would insert the morpholine ring into residues flanking the pocket and create steric clash. Using analogous chemistry, an amide coupling reaction was employed to furnish analog **4** (Scheme 2).

### Potency and Selectivity Analyses Enable Probe Nomination

Once prepared, compounds **2** and **4** were evaluated in the CDKL5 NanoBRET assay, which measures in-cell target engagement. As shown in Figure 3, compound **2** was found to have an IC_50_ value of 4.6 nM in the CDKL5 NanoBRET assay, while compound **4** was ∼1000× less active (IC_50_ = 4400 nM). Parent compound AT-7519 was also evaluated and demonstrated an IC_50_ value of 13 nM in the CDKL5 NanoBRET assay (Figure S1). These cellular target engagement values were confirmed using an orthogonal in vitro split-luciferase CDKL5 assay. The same trend was observed, with compound **2** demonstrating nearly 130-fold higher potency than compound **4** in this assay (Figure 3).

**Figure 3.**
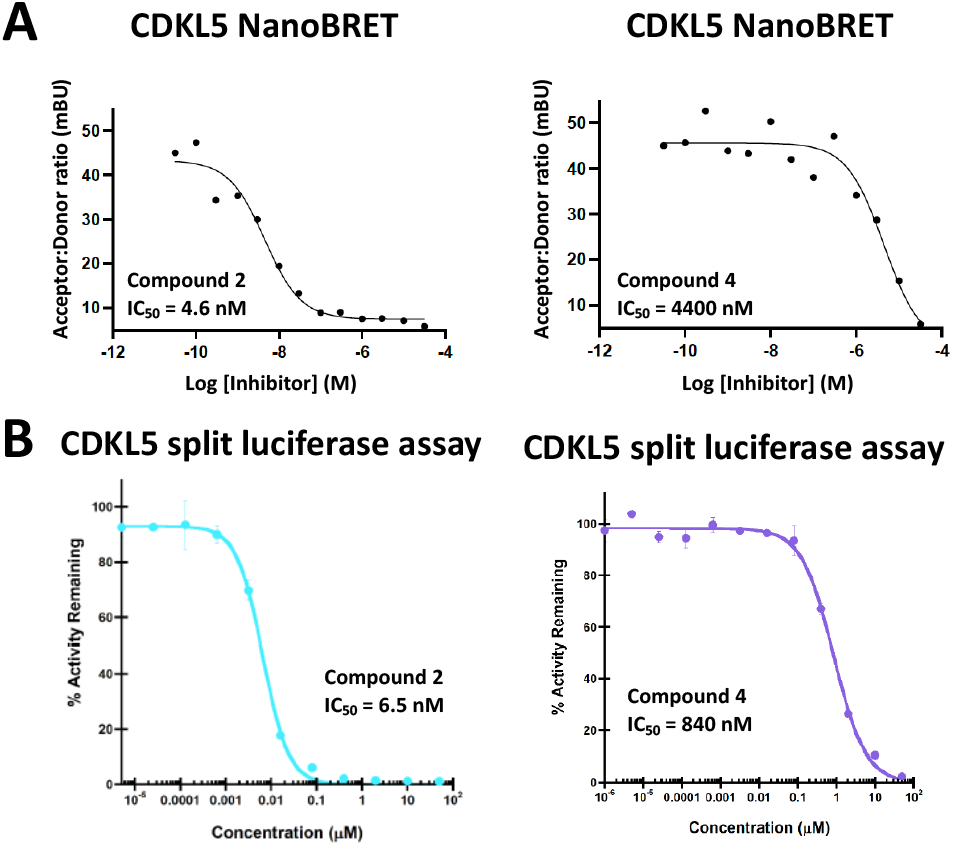
(A) CDKL5 cellular and (B) CDKL5 biochemical potency for compounds **2** and **4**.

The kinome-wide selectivity of AT-7519 has been characterized.^35^ After screening AT-7519 against 442 kinases at 10 μM, kinases that met affinity criteria were profiled in dose–response format to determine a quantitative dissociation constant (Kd) for each interaction. For 41 of the 442 kinases, a Kd value was calculated. As summarized in Figure S2, AT-7519 bound to 30 kinases with a Kd ≤1 μM. More than half of these kinases belong to the CDK or CDKL family.^35^

To determine whether the off-targets of AT-7519 were maintained for compounds **2** and **4**, we analyzed them at 1 μM in the Eurofins DiscoverX *scan*MAX panel. This platform assesses binding to 403 wild-type (WT) human as well as several mutant and non-human kinases, generating percent of control (PoC) values for each kinase evaluated.^35^ Compounds **2** and **4** proved to be exquisitely selective compounds, binding to only 4 and 5 kinases with PoC <10 at 1 μM, respectively. As shown in Figure 4, GSK3β and DYRK2 were common off-targets of both compounds. The PoC values for all CDKL family members included in the Eurofins DiscoverX *scan*MAX panel are included in Figure 4 as well. These data demonstrate the selectivity of these compounds for CDKL5 versus other CDKL kinases, which share a high degree of similarity.^12^ These binding assay results were corroborated when CDKL family selectivity was probed via thermal shift assays (Figure S4). The highest change in melting temperature (>6 °C) was observed for CDKL5 at all concentrations evaluated.

**Figure 4.**
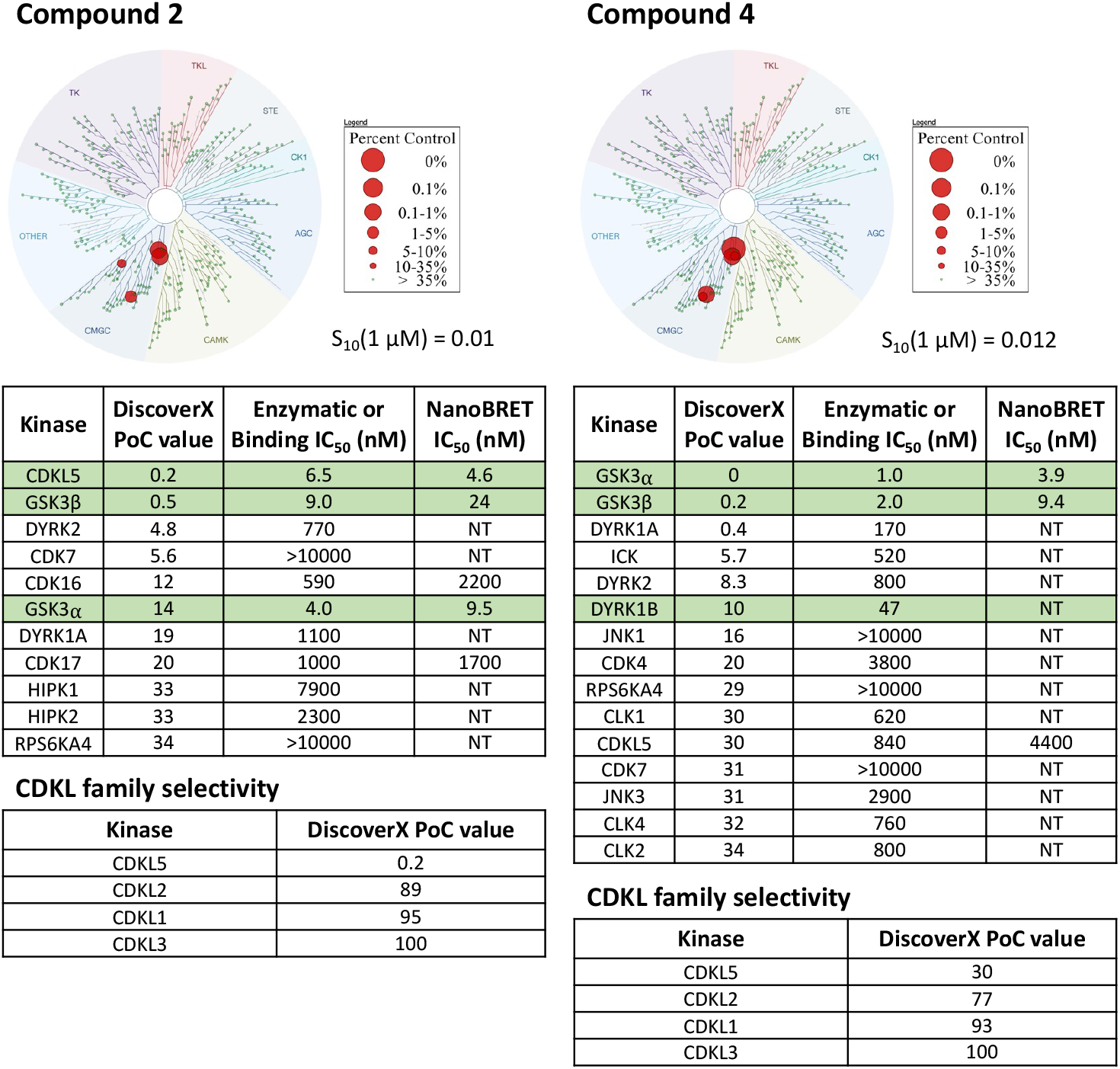
Selectivity data for compounds **2** and **4**. PoC: percent of control.

Corresponding enzymatic assays were run for all kinases with PoC <35 in the Eurofins DiscoverX *scan*MAX panel when compound **2** was profiled at 1 μM. Only GSK3α and β were potently inhibited within 30-fold of the CDKL5 enzymatic IC_50_ value for compound **2**. There is a 65-fold selectivity window between CDKL5 and the next most potently inhibited kinase (CDK16). Since GSK3α and β were confirmed to be potently inhibited, the corresponding GSK3α and β NanoBRET assays were run. The enzymatic potency translated to potent in cellular target engagement, with NanoBRET IC_50_ values <30 nM (Figure 5). Based on the pan-CDK activity of AT-7519, we also analyzed compound **2** in the CDK16 and CDK17 NanoBRET assays (Figure S3). The weaker enzymatic inhibition of these kinases translated to NanoBRET IC_50_ values between 1.5–2.3 μM. Together our data suggested that compound **2** is a potent, selective, and cell active CDKL5/GSK3 chemical probe.

**Figure 5.**
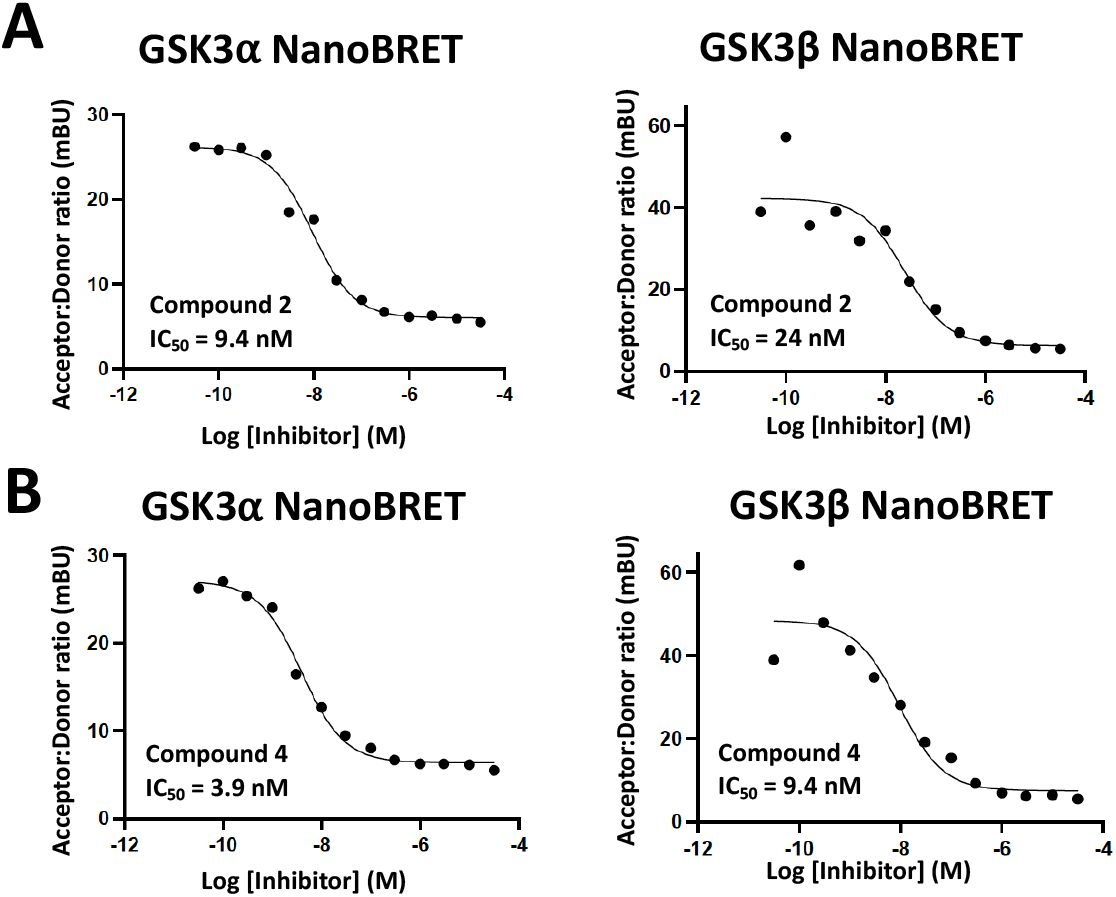
Cellular target engagement of GSK3α/β by compounds **2** and **4**.

Similarly, enzymatic assays were executed for all kinases with PoC <35 in the Eurofins DiscoverX *scan*MAX panel when analog **4** was screened at 1 μM. GSK3α and β were potently inhibited by compound **4**, a finding that was corroborated in the GSK3α and β NanoBRET assays (Figure 5). There is a 23-fold selectivity window between GSK3β and DYRK1B the next most potently inhibited kinase. It also lacks pan-CDK activity. Our profiling of compound **4** supports its selection as a suitable negative control to be paired with compound **2**. This assertion is based upon its lack of CDKL5 activity coupled with its narrow off-target inhibition profile. As compound **4** is a dual GSK3α/β chemical probe, its use in parallel with compound **2** will help illuminate activities resulting from CDKL5 versus GSK3α/β inhibition.

### Compound 2 Demonstrates Canonical ATP-Competitive Binding to CDKL5

As shown in Figures 6 and S6, compound **2** acts as a Type I kinase inhibitor and occupies the ATP pocket of CDKL5. The presence of a catalytic salt bridge between Lys42 and Glu60 suggests that CDKL5 is in an active conformation when this compound binds. Key interactions between compound **2** and CDKL5 include hydrogen bonding between hinge residues Glu90 and Val92 and the pyrazole N and NH. An additional hydrogen bond is observed between hinge residue Val92 and the 3-position amide NH. The piperidinyl nitrogen is also positioned suitably to form hydrogen bonds with Glu98 and the backbone carbonyl of Glu93. Favorable π-stacking was noted between Tyr24 and the difluorophenyl ring of compound **2**. Much of the compound assumes a planar overall geometry when bound, a finding that is partially influenced by constraints within the ATP binding pocket. Overall, the crystal structure of compound **2** is in agreement with the proposed binding mode of AT-7519 based on our earlier docking studies (Figure 1), validating our structure-driven hypotheses. Our co-crystal structure supports the idea that, for CDKL5 negative control compound **4**, the additional methylene inserted in the chain that connects to the tetrahydropyran precludes a hydrogen bond with Val92 in the hinge region of the ATP binding pocket. An unfavorable steric clash with the ATP site in the region of Val92 likely arises, which alters the binding orientation of this analog such that other key interactions are not maintained. If planarity of compound **4** cannot be achieved, then the constrained pocket is unlikely to tolerate its binding.

**Figure 6.**
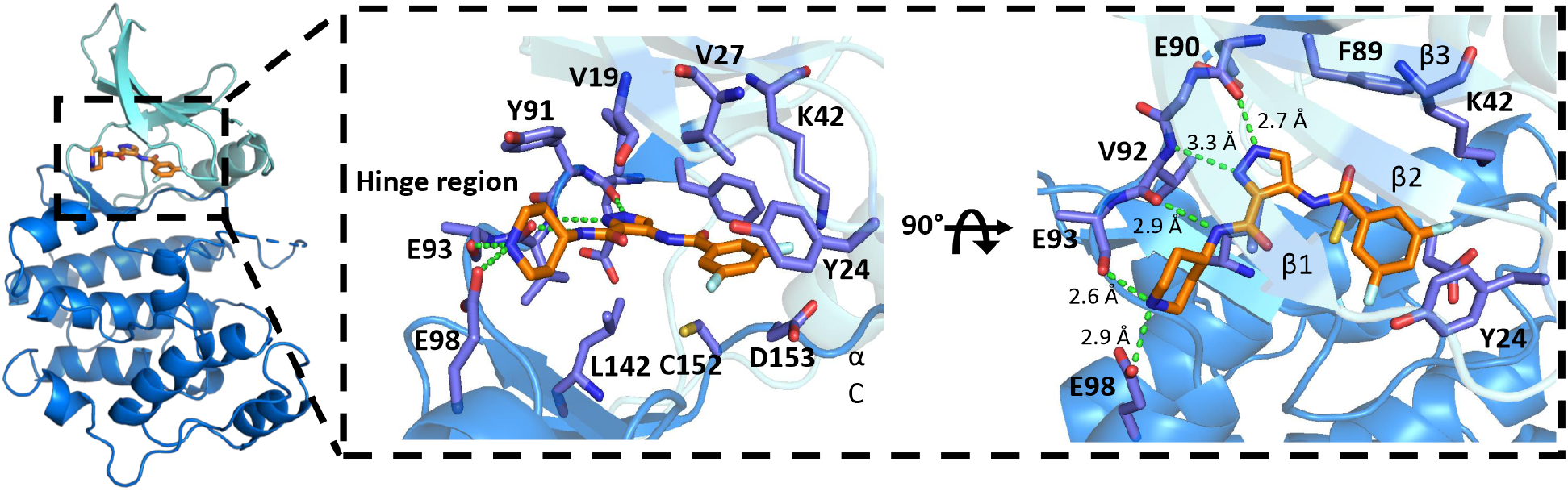
Overview of the CDKL5 co-crystal structure with compound **2** (PDB code: 8CIE). N and C terminal lobes colored in aquamarine and blue, respectively. Compound **2** binds to the ATP pocket between the two lobes and is shown as orange sticks. Hydrogen bonds are indicated as green dashed lines, including interactions between the hinge region (Glu90 and Val 92, colored purple) and the pyrazole shown in the cutout.

### CDKL5 Kinase Activity and Downstream Signaling are Inhibited by Chemical Probe

**2**. *In vitro* kinase assays were used to assess the impact of our chemical probe set on CDKL5 activity (Figure 7). Human WT and kinase dead (KD, CDKL5 K42R) recombinant proteins were prepared. When equal amounts were used in CDKL5 activity assays (Figure 7B), WT CDKL5 but not KD CDKL5 demonstrated kinase activity in the presence of ATP (50 μM, Figure 7A). Next, untreated WT CDKL5 was compared with WT CDKL5 treated with vehicle (DMSO) or 10 or 100 nM of a small molecule. Our chemical probe set (compounds **2** and **4**), AST-487, and lapatinib were included in this study. AST-487 represents a validated, but less selective inhibitor of CDKL5 (positive control),^12^ while lapatinib is a relatively selective kinase inhibitor that is not an inhibitor of CDKL5 (negative control). No significant difference was observed between treatments with DMSO, compound **4**, lapatinib and untreated CDKL5 at either concentration (Figure 7C,D). In contrast, compound **2** and AST-487 demonstrated robust, dose-dependent inhibition of CDKL5 activity (Figure 7C,D). These assays confirmed that compound **2** is a potent inhibitor of CDKL5 kinase activity.

**Figure 7.**
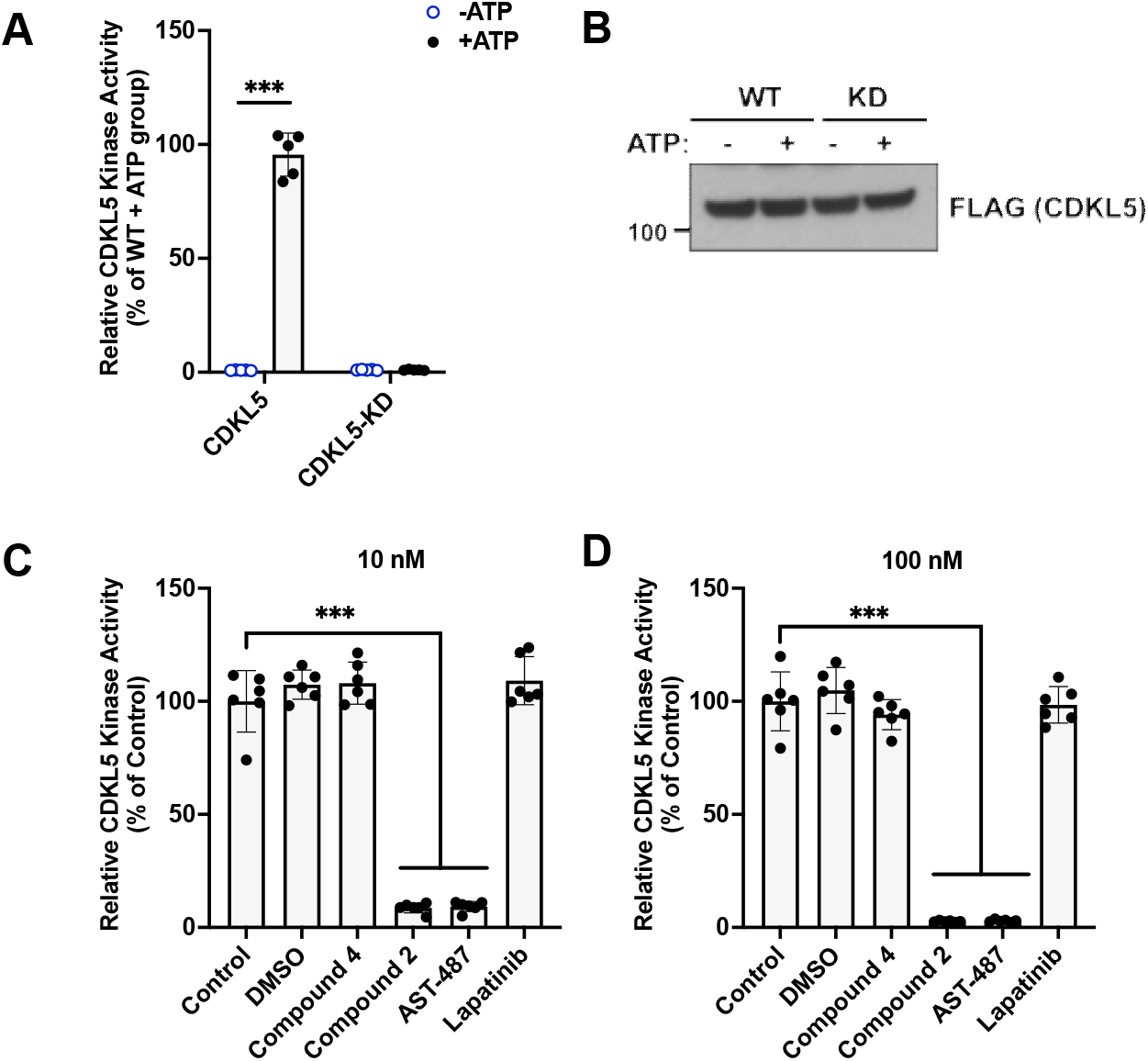
Compound **2** potently inhibits CDKL5 kinase activity. (A) Representative graph from a kinase assay showing that purified WT human CDKL5 retains kinase activity, while the kinase dead (KD, CDKL5 K42R) human protein was functionally inactive. n = 3 replicates. (B) Representative western blot showing equal levels of WT and KD proteins in the kinase assay experiments. (C-D) Purified WT human CDKL5 was used for kinase assays as described above in the presence of indicated compounds at 10 or 100 nM concentrations. n = 3 replicates. One-way ANOVA with Dunnett’s multiple comparison test was done. *** = p <0.0001. Non-significant comparisons are not illustrated.

CDKL5 as well as GSK3α and β are known to play essential roles in the brain. For this reason, we elected to study the impact of our chemical probe (**2**) and negative control (**4**) on specific signaling pathways in DIV15-16 cortical primary rat neurons. These neurons were treated with compounds **2** and **4** in dose–response format for 1 hour. As shown in Figures 8 and S5, we examined the response of CDKL5, EB2, phospho-EB2, phospho-GSK3α/β, β-catenin, and phospho-β-catenin to compound treatment. CDKL5 expression was not altered due to treatment with either compound. EB2, a microtubule-associated protein, is a known substrate of CDKL5.^25^ Phosphorylation of EB2 was inhibited in a dose-dependent manner by compound **2** and only at the highest concentration of compound **4**. The response to compound **2** at 5 nM and to compound **4** at 5 μM correlate well with their respective CDKL5 NanoBRET IC_50_ values of 4.6 nM and 4400 nM (Figure 3). This result also confirms that inhibition of CDKL5 by these compounds results in disruption of CDKL5 signaling in cells, albeit at 1000-fold different concentrations. Phosphorylation of β-catenin, a known substrate of GSK3α/β,^36^ was reduced in a dose-dependent manner by both compounds with a clear response in the 5 nM–50 μM range. Inhibition of β-catenin phosphorylation occurred within a concentration range that correlates well with the GSK3α/β NanoBRET IC_50_ values (3.9–24 nM) for compounds **2** and **4** (Figure 5). This result supports that both compounds inhibit the downstream signaling mediated by GSK3α/β in cells.

**Figure 8.**
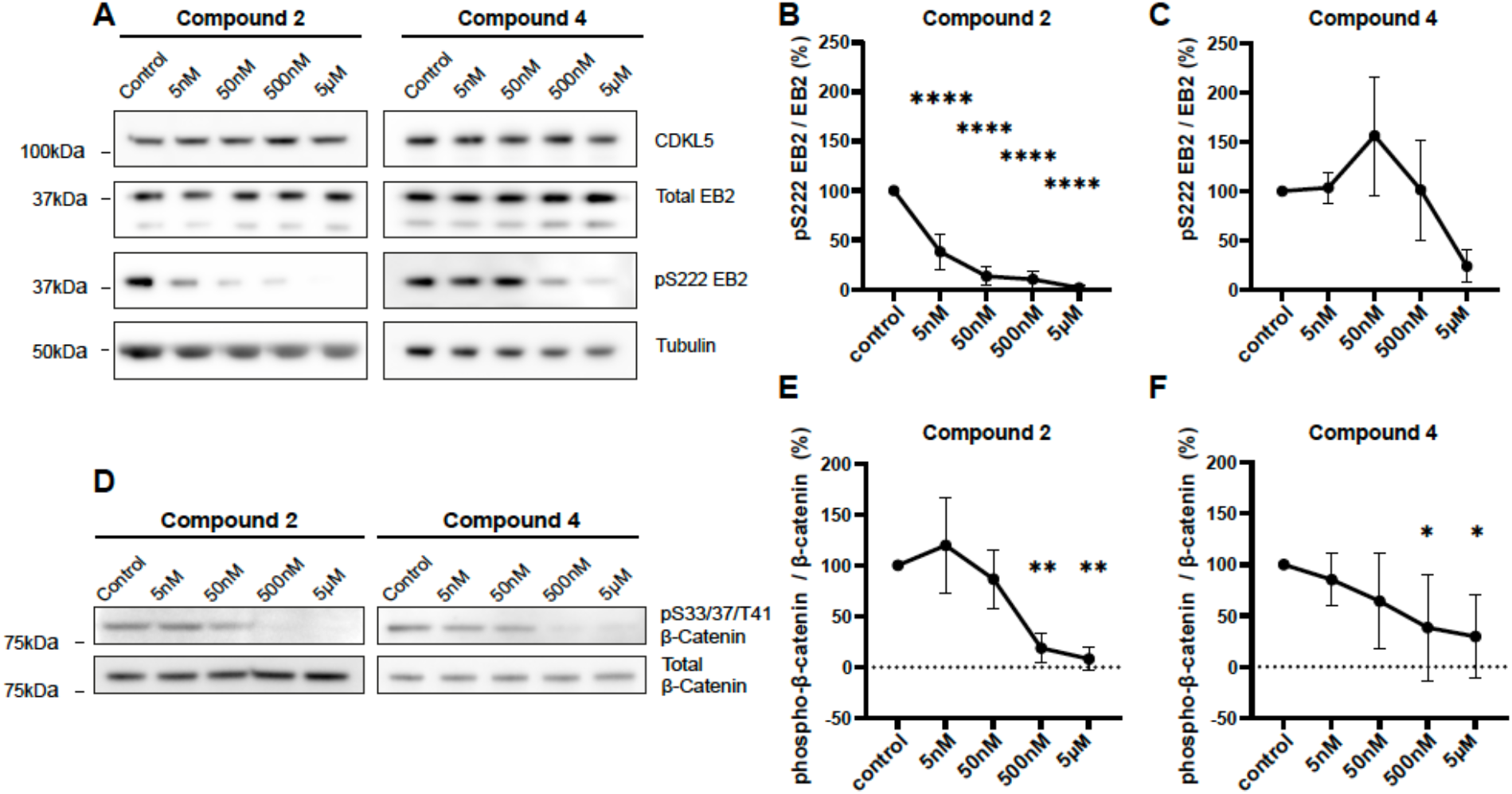
Analysis of phospho-EB2 and phospho-β-catenin expression following treatment with compound **2** or **4** confirms inhibition of downstream signaling mediated by CDKL5 and GSK3α/β. (A) Western blots showing expression of CDKL5, EB2 and tubulin, and level of S222 EB2 phosphorylation following 1 hour treatment of DIV11-14 rat primary cortical neurons with 5 nM, 50 nM, 500 nM or 5 μM of compound **2** or **4**. (B) Quantification of S222 EB2 phosphorylation following treatment of DIV11-14 rat primary cortical neurons with compound **2**. Two-way ANOVA with pairwise comparisons of each condition against the control condition. n = 3 replicates. (C) Quantification of S222 EB2 phosphorylation following treatment of DIV11-14 rat primary cortical neurons with compound **4**. Two-way ANOVA with pairwise comparisons of each condition against the control condition. n = 3 replicates. (D) Western blots showing expression of β-Catenin, and level of S33/37/T41 β-Catenin phosphorylation following 1 hour treatment of DIV11-14 rat primary cortical neurons with 5 nM, 50 nM, 500 nM or 5 μM of compound **2** or **4**. (E) Quantification of S33/37/T41 β-catenin phosphorylation following treatment of DIV11-14 rat primary cortical neurons with compound **2**. Two-way ANOVA with pairwise comparisons of each condition against the control condition. n = 3 replicates. (F) Quantification of S33/37/T41 β-catenin phosphorylation following treatment of DIV11-14 rat primary cortical neurons with compound **4**. Two-way ANOVA with pairwise comparisons of each condition against the control condition. n = 3 replicates. * = p ≤ 0.05, ** = p ≤ 0.01, *** = p ≤ 0.001, **** = p ≤ 0.0001. Non-significant (p > 0.05) comparisons are not illustrated.

### Chemical Probe Pair are Neuroprotective in Human Motor Neurons

Amyotrophic lateral sclerosis (ALS) is characterized by the widespread and rapid degeneration of motor neurons in the brain and spinal cord. Like many other neurodegenerative disorders, disease-causing mutations in ALS lead to the accumulation of misfolded proteins, triggering endoplasmic reticulum (ER) stress and activating the unfolded protein response (UPR).^15, 16^ The UPR slows protein synthesis and promotes refolding or clearance of misfolded proteins to protect neurons and support their survival.^16^ If recovery is not achieved, however, the UPR will drive apoptotic signaling to kill ER-stressed cells.^16^ Markers of ER stress are among the earliest pathological features to appear in animal models of ALS.^37^ Motor neurons are significantly more sensitive than other spinal neuronal subtypes to pharmacologically induced ER stress,^38^ which may help explain their enhanced vulnerability in ALS. At the same time, compounds that target kinases have proven to protect motor neurons from ER stress.^38^ Thus, preventing ER stress-mediated neurodegeneration could be a promising therapeutic approach for patients with these diseases.

As part of their neuroprotective profile, GSK3 inhibitors reproducibly protect cells from ER stress-induced apoptosis.^16^ Less is known about the impact of CDKL5 inhibition on mediating ER stress pathways. To compare the pro-survival phenotype elicited by CDKL5 plus GSK3α/β inhibition versus GSK3α/β inhibition alone in response to ER stress, we treated differentiated human motor neurons derived from human stem cells with compounds **2** and **4**. Cells were treated with AT-7519 in parallel. ER stress was pharmacologically induced via treatment with cyclopiazonic acid, a mycotoxin that inhibits calcium transport within the ER.^39^ Human motor neurons were selected as a neuronal subtype that is degenerated in ALS.^40^

All three compounds were found to be neuroprotective (Figure 9). At low concentrations (<200 nM), which align with their IC_50_ values in the CDKL5 and GSK3α/β NanoBRET assays, compounds **2** and **4** better promoted motor neuron survival than AT-7519. We know less about the selectivity of these compounds at concentrations >1 μM but suggest that several additional kinases will be inhibited. Since only subtle differences were observed in response to compound **2** versus **4**, we conclude that CDKL5 inhibition is not detrimental to motor neuron survival. This contrasts with reports that CDKL5 inhibition without GSK3α/β inhibition is harmful. Specifically, that loss of CDKL5 in mice induces abnormal activity of GSK3β, leading to insufficient neuronal maturation and increased apoptosis.^30^ This GSK3β activation can be corrected via treatment with GSK3β inhibitors.^19, 20^ Compound **4** is a potent and selective GSK3α/β inhibitor that shows promise due to its neuroprotective nature. These data demonstrate that the compounds in our probe set are not toxic to human iPSC-derived motor neurons and could be useful for studies with a neurological focus.

**Figure 9.**
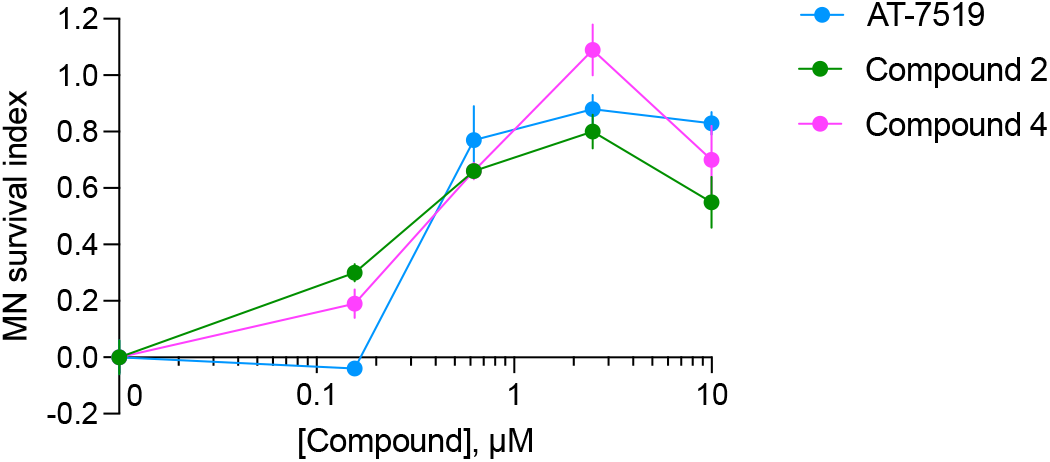
Compounds **2** and **4** promote survival of motor neurons in response to ER stress. Error bars represent standard error of the mean (SEM) calculated using two-way ANOVA. MN: motor neuron.

## CONCLUSION

In summary, we designed and synthesized a series of AT-7519 analogs from which we identified a CDKL5/GSK3 chemical probe (**2**). This compound demonstrates potent inhibition of CDKL5 and GSK3α/β and retains high affinity for these kinases in cells. Making subtle structural changes to afford analog **4** abolished its ability to bind to CDKL5 without jeopardizing its affinity for GSK3α/β. Our co-crystal structure of chemical probe **2** bound to CDKL5 suggests that the ATP site of CDKL5 would likely not accommodate compound **4**. This is hypothesized to be due to an unfavorable steric clash resulting from its longer side chain versus compound **2**. Kinome-wide profiling of both members of the chemical probe set confirmed excellent broad kinome selectivity. Our studies in human motor neurons highlight the neuroprotective nature of our chemical probe set. Compounds **2** and **4** promoted survival of motor neurons submitted to endoplasmic reticulum dysfunction, which is a hallmark of neurodegenerative diseases like ALS. Based on its potency, selectivity, and activity in cells, we suggest analog **2** as an optimal CDKL5/GSK3 chemical probe and, due to its lower CDKL5 potency and selectivity, analog **4** to be a suitable negative control to be distributed and used alongside it. In addition, we suggest that their neuroprotective nature makes them suitable candidates for exploratory cell-based models of CNS diseases.

## METHODS

### Chemical Synthesis

#### General Information

Reagents were obtained from trusted commercial sources and used without further analytical characterization. Solvent was removed via a rotary evaporator under reduced pressure and thin layer chromatography in tandem with LC–MS was used to monitor the reaction progress. The following abbreviations are used in schemes and/or experimental procedures: mmol (millimoles), mg (milligrams), equiv (equivalent(s)), and h (hours). ^1^H NMR and other microanalytical data were collected for final compounds to confirm their identity and assess their purity. ^1^H and ^13^C NMR spectra were collected in methanol-*d*_*4*_ using Bruker spectrometers with the magnet strength indicated in each corresponding line listing. Peak positions are noted in parts per million (ppm) and calibrated versus the shift of methanol-*d*_*4*_; coupling constants (*J* values) are listed in hertz (Hz); and multiplicities are included as follows: singlet (s), doublet (d), doublet of doublets (dd), doublets of doublets (ddd), doublet of triplet of doublets (dtd), triplet (t), triplet of doublets/triplets (td/tt), triplets of triplets (ttt), and multiplet (m).

#### 4-(3,5-difluorobenzamido)-N-(piperidin-4-yl)-1H-pyrazole-3-carboxamide (2)

A solution of *tert*-butyl 4-(4-amino-1*H*-pyrazole-3-carboxamido)piperidine-1-carboxylate^34^ (75.0 mg, 0.242 mmol, 1.0 equiv), 3,5-difluorobenzoic acid (38.3 mg, 0.242 mmol, 1.0 equiv), and DIPEA (160 μL, 0.970 mmol, 4.0 equiv) in THF (5.0 mL) was cooled to 0 °C and treated dropwise with *n*-propanephosphonic acid anhydride (T_3_P, 50% solution in ethyl acetate, 231 μL, 0.364 mmol, 1.5 equiv). The reaction mixture was then stirred for 1 h at room temperature (25 °C). Upon completion, solvent was removed under reduced pressure and the residue was dissolved in ethyl acetate. The organic phase was washed with water and the product was extracted with ethyl acetate (2 × 5 mL). The organic solution was dried over anhydrous Na_2_SO_4_ and concentrated *in vacuo*. The residue was purified with column chromatography (SiO_2_, 10–100% ethyl acetate in hexane) to yield *tert*-butyl 4-(4-(3,5-difluorobenzamido)-1*H*-pyrazole-3-carboxamido)piperidine-1-carboxylate (51.0 mg, 47% yield), which was carried on without further characterization.

A solution of *tert*-butyl 4-(4-(3,5-difluorobenzamido)-1*H*-pyrazole-3-carboxamido)piperidine-1-carboxylate (43.0 mg, 0.096 mmol, 1 equiv) in dioxane (1 mL) was cooled to 0 °C and treated dropwise with hydrogen chloride solution (4.0 M in dioxane, 3.0 mL). The reaction mixture was then stirred for 1 h at room temperature (25 °C). Upon completion, solvent was removed under reduced pressure. The residue was purified with column chromatography (SiO_2_, 10–100% ethyl acetate in hexane) to yield compound **2** (29.0 mg, 79% yield). ^1^H NMR (850 MHz, methanol-*d*_4_) δ 8.32 (s, 1H), 7.54 – 7.49 (m, 2H), 7.26 (tt, *J* = 8.8, 2.3 Hz, 1H), 4.22 (tt, *J* = 10.9, 4.1 Hz, 1H), 3.52 – 3.44 (m, 2H), 3.18 (td, *J* = 12.8, 3.1 Hz, 2H), 2.22 (dd, *J* = 14.3, 3.2 Hz, 2H), 1.96 – 1.87 (m, 2H). ^13^C NMR (214 MHz, Methanol-*d*_4_) δ 165.58, 164.70 (dd, *J* = 249.4, 12.4 Hz), 162.96 (t, *J* = 2.7 Hz), 138.48 (t, *J* = 8.5 Hz), 134.32, 123.94, 121.99, 111.34 (dd, *J* = 22.3, 4.7 Hz), 108.33 (t, *J* = 25.8 Hz), 45.33, 44.29, 29.51. HRMS: calcd for C_16_H_18_F_2_N_5_O_2_ [M + H]^+^ m/z, 350.1429; found m/z, 350.1421.

#### 4-(2,6-difluorobenzamido)-N-((tetrahydro-2H-pyran-4-yl)methyl)-1H-pyrazole-3-carboxamide (**4**)

A solution of 4-(2,6-difluorobenzamido)-1*H*-pyrazole-3-carboxylic acid^34^ (42.0 mg, 0.157 mmol, 1.0 equiv), 4-aminomethyltetrahydropyran (21.3 μL, 0.189 mmol, 1.2 equiv), and DIPEA (77.9 μL, 0.472 mmol, 3.0 equiv) in THF (5.0 mL) was cooled to 0 °C and treated dropwise with *n*-propanephosphonic acid anhydride (T_3_P, 50% solution in ethyl acetate, 150 μL, 0.236 mmol, 1.5 equiv). The reaction mixture was then stirred for 1 h at room temperature (25 °C). Upon completion, solvent was removed under reduced pressure and the residue was dissolved in ethyl acetate. The organic phase was washed with water and the product was extracted with ethyl acetate (2 × 5.0 mL). The organic solution was dried over anhydrous Na_2_SO_4_ and concentrated *in vacuo*. The residue was purified with column chromatography (SiO_2_, 10–100% ethyl acetate in hexane) to yield compound **4** (15.5 mg, 27% yield). ^1^H NMR (850 MHz, methanol-*d*_4_) δ 8.34 (s, 1H), 7.58 (tt, *J* = 8.4, 6.3 Hz, 1H), 7.14 (t, *J* = 8.3 Hz, 2H), 3.94 (ddd, *J* = 11.8, 4.7, 2.0 Hz, 2H), 3.40 (td, *J* = 11.8, 2.2 Hz, 2H), 3.25 (d, *J* = 7.0 Hz, 2H), 1.87 (ttt, *J* = 11.0, 7.1, 3.8 Hz, 1H), 1.68 (ddd, *J* = 13.1, 4.0, 2.0 Hz, 2H), 1.33 (dtd, *J* = 13.5, 11.9, 4.6 Hz, 2H). ^13^C NMR (214 MHz, methanol-*d*_4_) δ 165.78, 161.49 (dd, *J* = 252.0, 6.5 Hz), 158.91, 134.42, 134.17 (t, *J* = 10.5 Hz), 123.14, 122.15, 114.67 (t, *J* = 19.7 Hz), 113.33 (dd, *J* = 22.0, 3.7 Hz), 68.72, 45.41, 36.66, 31.81. HRMS: calcd for C_17_H_19_F_2_N_4_O_3_ [M + H]^+^ m/z, 365.1425; found m/z, 365.1419.

### Kinome-Wide Selectivity Analyses

The *scan*MAX assay platform (Eurofins DiscoverX Corporation) was used to assess the broad selectivity of compounds **2** and **4** at 1 μM. These compounds were screened against 403 wild-type (WT) human kinases and percent of control (PoC) values were generated, allowing for calculation of selectivity scores (S_10_(1 μM)).^35^ Selectivity scores, the number of WT human kinases in the *scan*MAX panel with PoC <10, and PoC values corresponding with all CDKL family member kinases within the panel are included in Figure 4. Kinome tree diagrams, also in Figure 4, were generated based on this binding data for compounds **2** and **4**. A kinome tree diagram for AT-7519 was generated based on published data and included in Figure S2.^35^

### Enzymatic and Biochemical Assays

IC_50_ determinations against CDKL5 were executed using the *KinaseSeeker* homogenous competition binding assay at Luceome Biotechnologies, LLC.^41^ Briefly, this luminescence-based assay depends on displacement of an active site dependent probe by an inhibitor. Compounds were tested in dose–response (12-pt curve) format in duplicate. Representative curves are included in Figure 3 and corresponding IC_50_ values are embedded in tables within Figure 4.

Eurofins enzymatic radiometric assays were run at the K_m_ value for ATP to generate dose– response (9-pt) curves for all kinases listed in Figure 4 with the exception of CDKL5 and DYRK1B. The protein constructs, substrate and controls employed, additional proteins added, and detailed assay protocol for these assays can be found on the Eurofins website: https://www.eurofinsdiscoveryservices.com.

The radiometric HotSpot kinase assay for DYRK1B was executed at Reaction Biology Corp. (RBC) at the K_m_ value for ATP in dose–response (10-pt curve) format. The IC_50_ value generated is included in the embedded table within Figure 4 corresponding with compound **4**. Assay details for this assay can be accessed via the RBC website: https://www.reactionbiology.com/list-kinase-targets-us-facility.

### CDKL5 Protein Expression and Purification

Baculoviral expression of human CDKL5 (UniProt O76039; residues 1–303; T169D and Y171E) was performed in Sf9 cells as described previously.^12^ Cells were harvested 72 hr post-infection and resuspended in 30 mL binding buffer (50 mM HEPES pH 7.5, 500 mM NaCl, 5% glycerol) supplemented with 0.01% Triton X-100, 1 mM TCEP and protease inhibitors. Cells were lysed by sonication. Polyethylenimine was added to a final concentration of 0.5% to precipitate DNA and 21 μL of 25 mM compound **2** was then added to stabilize the CDKL5 protein during purification. The cell lysate was clarified by centrifugation and proteins purified by nickel-affinity and size exclusion chromatography. 10 mM arginine/glutamate mix (final concentration) was added before protein concentration. Tobacco etch virus protease A (TEV) was used to proteolytically cleave the polyhistidine tag overnight at 4°C. **Crystallization**. Crystals were grown at 20°C in a precipitant containing 0.1 M tri-sodium citrate, 1.1 M lithium sulfate, and 0.4 M ammonium sulfate. 15% glucose was added as a cryoprotectant for crystal mounting before vitrification in liquid nitrogen.

### Diffraction Data Collection, Structure Solution and Refinement

Diffraction data were collected at Diamond Light Source on beamline i04 to a resolution of 2.2 Å. Data were processed with the Xia2 pipeline^42^, using DIALS software.^43^ PDB 4BGQ was used as a search model for molecular replacement in PHENIX.^44^ The initial model was improved by rounds of manual building in Coot^45^ and refined by PHENIX.^44^ Data collection and refinement statistics are provided in Table S1.

### Thermal Shift Assays

CDKL1, CDKL2, CDKL3, and CDKL5 were generated using the construct boundaries previously used in the generation of 4AGU, 4AAA, 3ZDU, and 4BGQ, respectively. The CDKL1, CDKL2, CDKL3, or CDLK5 kinase domain at 4 μM in 10 mM HEPES-NaOH pH 7.4 and 500 mM NaCl was incubated with the inhibitors at different concentrations (12.5, 25, or 50 μM) in the presence of 5× SyPRO orange dye (Invitrogen). A Real-Time PCR Mx3005p machine (Stratagene) was used to record fluorescence. A previously described protocol was followed to execute the Tm shift assays and evaluate for melting temperatures.^46^

### Cell Culture

Human embryonic kidney (HEK293) cells were obtained from ATCC and cultured in Dulbecco’s Modified Eagle’s medium (DMEM, Gibco) supplemented with 10% (v/v) fetal bovine serum (FBS, Corning). Culture conditions included incubation in 5% CO_2_ at 37°C and passaging every 72 hours with trypsin (Gibco). Cells were never allowed to reach confluency.

### Rat primary culture

Pregnant Long Evans rats were ordered from Jackson labs. E18.5 rat embryos were removed from the uterus and the brains were taken out. Cortices were dissected out, pooled from multiple animals, and washed three times with Hank’s Balanced Salt Solution (HBSS). An incubation with 0.25 % trypsin for 15 minutes at 37 °C was followed by four times washing with HBSS. Cells were dissociated and then counted using a hemocytometer. Neurons were plated on 12-well culture plates at a density of 300,000 cells per well. The wells were coated with 0.1 M borate buffer containing 60 μg/mL poly-D-lysine and 2.5 μg/mL laminin, and placed in the incubator overnight. Neurons were plated with minimum essential medium (MEM) containing 10% fetal bovine serum (FBS), 0.5% dextrose, 0.11 mg/mL sodium pyruvate, 2 mM glutamine, and penicillin/streptomycin. After 4 hours, cultures were transferred to neurobasal medium containing 1 mL of B27 (Gibco), 0.5 mM glutamax, 0.5 mM glutamine, 12.5 μM glutamate, and penicillin/streptomycin. Primary neuronal cultures were kept at 37 °C and 5% CO_2_. Every 3–4 days, 20–30% of the maintenance media was refreshed. Between DIV11-14, neurons were treated with 5 nM, 50 nM, 500 nM and 5 μM of compound **2** or compound **4** for 1 hour. The compounds were added directly to the media and the plates were placed at 37 °C for the time of the treatment. DMSO was added to the well for the control condition.

### Human Motor Neuron Differentiation and Maintenance

Human motor neurons were derived from the wild-type male iPS cell line, NCRM-1 (RRID:CVCL_1E71), carrying a VAChT:TdTomato reporter, as previously described.^47^ Induced pluripotent stem cells (iPSCs) were maintained at 37°C in humidified incubators on irradiated CF-1 mouse embryonic fibroblast (MEF) feeder layers (Thermo Fisher) in serum-free media (DMEM/F12, Life Technologies) supplemented with the following: Glutamax (1%, Life Technologies), knockout serum replacement (20%, Life Technologies), non-essential amino acids (1%, Millipore), human recombinant fibroblast growth factor 2 (FGF2, 20 ng/mL, Life Technologies), and beta-mercaptoethanol (0.1%, Sigma).^38^ iPSCs were then differentiated into motor neurons according to published protocols.^38, 48^ Briefly, iPSCs were dissociated into single cells via Accutase (Thermo Fisher) and differentiated into motor neurons as embryoid bodies cultured in suspension at 37°C over a time course of 16 days. Motor neuron differentiation was executed in N2B27 media comprised of: Advanced DMEM/F12 and Neurobasal media (1:1 mixture, Life Technologies), beta-mercaptoethanol (0.1%), Glutamax (1%), ascorbic acid (10 μM, Sigma), B-27 (2%, Thermo Fisher), and N-2 (1%, Thermo Fisher). Differentiation media was supplemented as needed over the 16-day period.^38^

### NanoBRET Assays

Constructs for NanoBRET measurements of CDKL5 (NLuc-CDKL5), GSK3α (NLuc-GSK3α), GSK3β (NLuc-GSK3β), CDK16 (CDK16-NLuc), and CDK17 (CDK17-NLuc), included in Figures 3–5, S1, and S3, were kindly provided by Promega. N-terminal NLuc orientations were used for CDKL5, GSK3α, and GSK3β, while C-terminal NLuc constructs were employed for CDK16 and CDK17. NanoBRET assays were executed in dose–response (5-pt or 12-pt curves) format as previously reported.^49^ Assays were carried as described by the manufacturer using 0.31 μM of tracer K11 for CDKL5, 0.13 μM of tracer K8 for GSK3α, 0.063 μM of tracer K8 for GSK3β, 0.5 μM of tracer K10 for CDK16, and 0.5 μM of tracer K10 for CDK17. Cyclin Y was added to the CDK16 and CDK17 NanoBRET assays in accordance with the manufacturer’s instructions.^50^

### CDKL5 Enzymatic Assays

Methods used for these assays have been described.^28^ Briefly, human FLAG-tagged CDKL5 WT or kinase dead (KD, CDKL5 K42R) constructs were subcloned into pT7CFE1-CHis plasmid (Thermo Fisher). These constructs were then used for *in vitro* translation using a HeLa cell lysate-based Kit (1-Step Human Coupled IVT Kit—DNA, 88881, Life Technologies). The *in vitro*-translated proteins were then purified using His Pur cobalt spin columns (Thermo Scientific). For *in vitro* kinase assays, recombinant CDKL5 and myelin basic protein (Active Motif, 31314) as a substrate were incubated in a kinase buffer (Cell Signaling, 9802) supplemented with or without 50 μM adenosine 5′-triphosphate (ATP) at 30 °C for 30 minutes followed by kinase assays using ADP-Glo Kinase Assay kit (Promega). Data were analyzed using GraphPad Prism 9. Each assay was run in triplicate (n = 3) and mean values are graphed in Figure 7. All error bars in Figure 7 are standard deviation (SD). Statistical analysis was done using one-way ANOVA with Dunnett’s multiple comparisons test. Statistical methods and p-values are mentioned in the Figure 7 legend. Thresholds for significance were placed at ***p<0.0001. Non-significant statistics were not indicated.

### Western Blots

After treatment, neuronal cultures were lysed in 300 μL of 1X sample buffer (Invitrogen) containing 0.1 M DTT. Lysates were briefly sonicated twice and denatured at 70 °C for 10 minutes. The samples were centrifuged at 13,300 rpm for 10 minutes and ran on NuPage 4-12% Bis-Tris polyacrylamide gels (Invitrogen). Proteins were transferred onto a Immobilon PVDF membrane (Millipore), which was then blocked in 4% milk in tris-buffered saline containing 0.1% Tween-20 (TBST) for 30 minutes. Primary antibodies were incubated at 4 °C overnight, and HRP-conjugated secondary antibodies at RT for two hours. The following primary antibodies were used: rabbit anti-CDKL5 (1:1,000; Atlas HPA002847), rabbit anti-pS222 EB2 (1:2,000, internal^25^), rat anti-EB2 (1:2,000; Abcam ab45767), mouse anti-tubulin (1:100,000; Sigma T9026), rabbit anti-β-catenin (1:1,000; Cell Signaling 9562), and rabbit anti phospho-β-catenin (1:250, Cell Signaling 9561). The following secondary antibodies were used at a concentration of 1:10,000: HRP-conjugated anti-rabbit (Jackson 711-035-152), HRP-conjugated anti-mouse (Jackson 715-035-151) and HRP-conjugated anti-rat (Jackson 712-035-153). The membrane was developed using ECL reagent (Cytiva) and was visualized with an Amersham Imager 600 (GE Healthcare). Quantification of Western blots was manually performed using Image Studio Lite Software (version 5.2). EB2 phosphorylation was measured relative to total EB2 and β-catenin phosphorylation to total β-catenin. The other proteins were normalized to tubulin, if not indicated otherwise.

Data were analyzed using GraphPad Prism 9. Exact values of n and statistical methods are mentioned in the figure legends. Each concentration was compared to the control using a Kruskal-Wallis test. n = 6 for each concentration. A p-value higher than 0.05 was not considered as statistically significant. Thresholds for significance were placed at *p ≤0.05, **p ≤0.01, ***p ≤0.001 and ****p ≤0.0001. All error bars in the figures are standard deviation (SD). Errors bars and non-significant statistics were not indicated.

### Neuroprotection Assay

On day 16 of differentiation, embryoid bodies were detached using trypsin (0.05%, Life Technologies) coupled with mechanical trituration. Dissociated motor neurons were then plated at 2000 cells/well in 96-well plates (Greiner) coated with mouse laminin (3 μg/mL, Thermo Fisher) and poly-ornithine (100 μg/mL, Sigma). Once plated, cells were maintained at 37°C in serum-free neurobasal media (Life Technologies) supplemented with a 1:1 cocktail of uridine and fluorodeoxyuridine as anti-mitotics (1 μM, Sigma), B-27 (2%), and N-2 (1%), plus 10 ng/mL of GDNF, BDNF, IGF-1 (R&D Systems), and CNTF (R&D Systems).^38^ Next, 48 hours after plating, motor neurons were treated simultaneously with cyclopiazonic acid (CPA) to induce ER stress and with the test compounds. Test compounds were added to three replicate wells in 4-fold dilutions from 0.16–10 μM. Motor neurons were co-incubated with CPA and test compounds for 72 hours. Live cells were then visualized using CellTrace Calcein AM and whole well images were acquired using a Plate RunnerHD system (Trophos). Cells were counted using Metamorph software (Molecular Devices). All cell counts are expressed as a percentage of surviving vehicle (DMSO)-treated cells. SEM was calculated using two-way ANOVA and plotted as error bars in Figure 8.

## Supporting information

Combined Supplemental Data

## ASSOCIATED CONTENT

### Supporting Information

The Supporting Information is available free of charge at https://pubs.acs.org/doi/TBD.

NanoBRET curves and NMR analysis of target compounds **2** and **4** and selectivity diagram for AT-7519 (PDF)

## AUTHOR INFORMATION

Complete contact information is available at: https://pubs.acs.org/TBD.

### Author Contributions

^⊥^H.W.O. and Y.L. contributed equally to this work. The manuscript was written and edited via contributions of all authors. All authors have approved of the final version of the manuscript.

### Notes

The authors declare no competing financial interests. The crystallographic coordinates of the CDKL5 co-structure have been deposited in the Protein Data Bank as 8CIE.

## ACKNOWLEDGMENTS

NanoBRET constructs for CDKL5, GSK3α, GSK3β, CDK16, and CDK17 were kindly provided by Promega. The TREE*spot* kinase interaction mapping software was employed in preparation of the kinome trees in Figures 4 and S2: http://treespot.discoverx.com. The authors would like to thank Diamond Light Source for beamtime (proposal mx28172), as well as the staff of beamline i04 for assistance with crystal testing and data collection.

## ABBREVIATIONS

AKT: protein kinase B
Amph1: amphiphysin
ANOVA: analysis of variance
ARHGEF2: Rho/Rac guanine nucleotide exchange factor 2
CDK: cyclin dependent kinase
CDKL: cyclin-dependent kinase-like
CEP131: centrosomal protein 131
DIPEA: N,N-diisopropylethyl amine
DLG5: Discs large MAGUK scaffold protein 5
Dnmt1: DNA methyltransferase
DYRK2: dual specificity tyrosine phosphorylation regulated kinase 2
EB2: microtubule end-binding protein 2
ELOA: elongin A
Glu: glutamic acid
H2B: histone H2B
HCl: hydrochloric acid
HDAC4: histone deacetylase 4
HIPK2: homeodomain interacting protein kinase 2
iPSC: induced pluripotent stem cell
Lys: lysine
MAP1S: microtubule associated protein 1S
MeCP2: Methyl-CpG-binding-protein
NGL-1: netrin-G ligand-1
PDK1: 3-phosphoinositide-dependent kinase 1
Rac1: Ras-related C3 botulinum toxin substrate 1
RT: room temperature
Sox9: SRY-box transcription factor 9
THF: tetrahydrofuran
Thr: threonine
Tyr: tyrosine
Val: valine

