## Supplementary material for "A Potent and Selective CDKL5/GSK3 Chemical Probe is Neuroprotective": Combined Supplemental Data

### Table of Contents

Figure S1 S2
Figure S2 S2
Figure S3 S3
Figure S4 S3
Figure S5 S4
Table S1 S5
Spectra for all compounds S6–S7

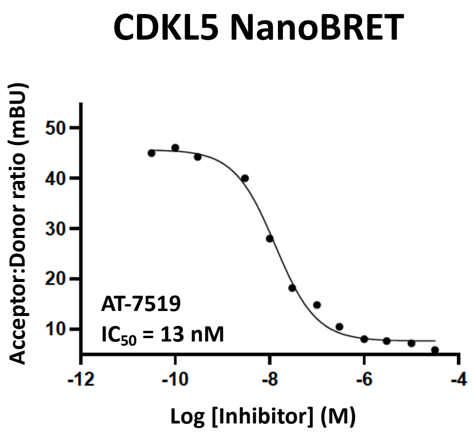


**Figure S1.** CDKL5 cellular target engagement by AT-7519.

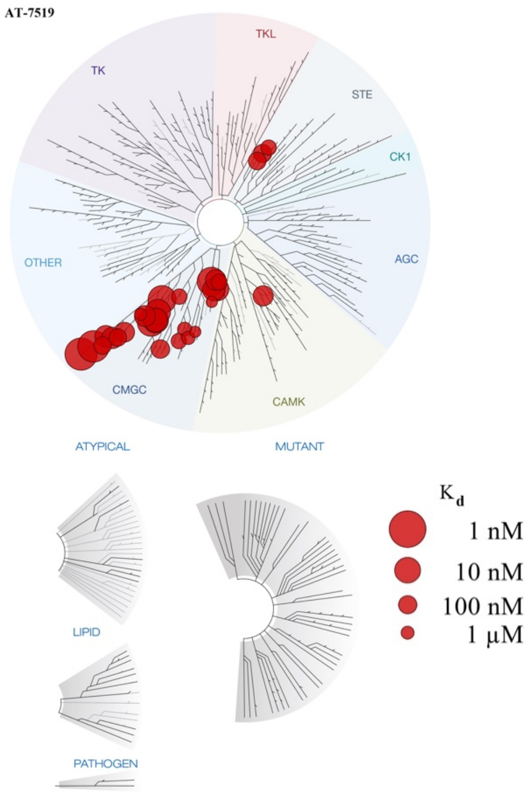


**Figure S2.** Kinome tree showing kinases that bind with a Kd value ≤1 μM when AT-7519 was profiled against 442 kinases.

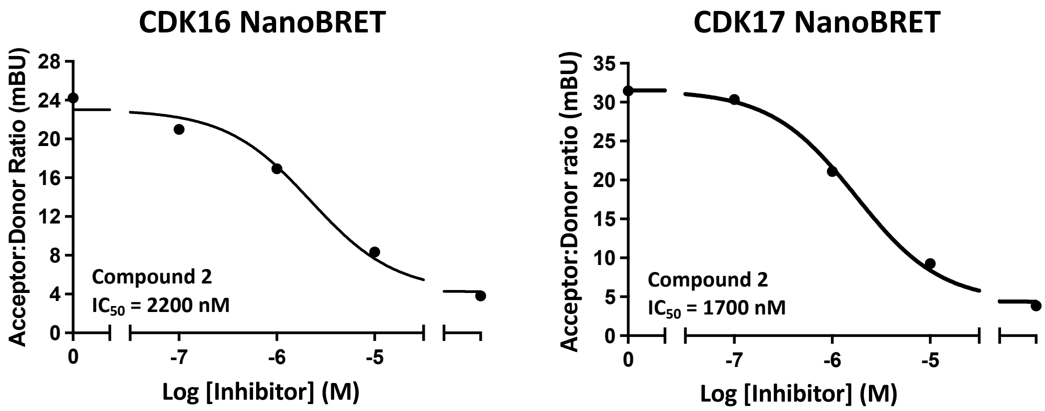


**Figure S3.** CDK16 and CDK17 cellular target engagement by compound **2**.

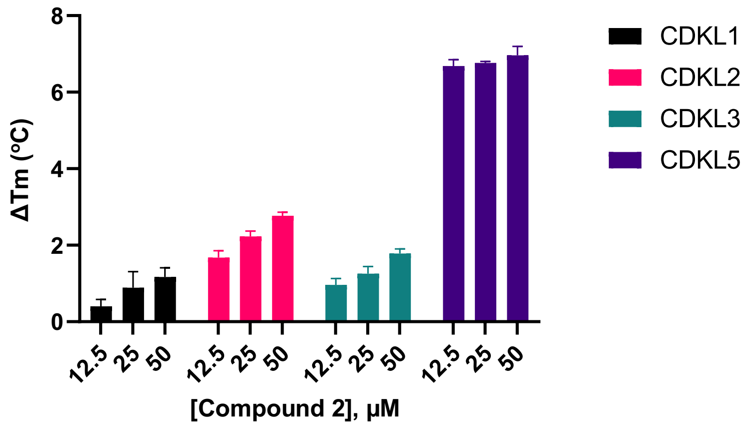


**Figure S4.** CDKL family selectivity evaluation via thermal shift assays.

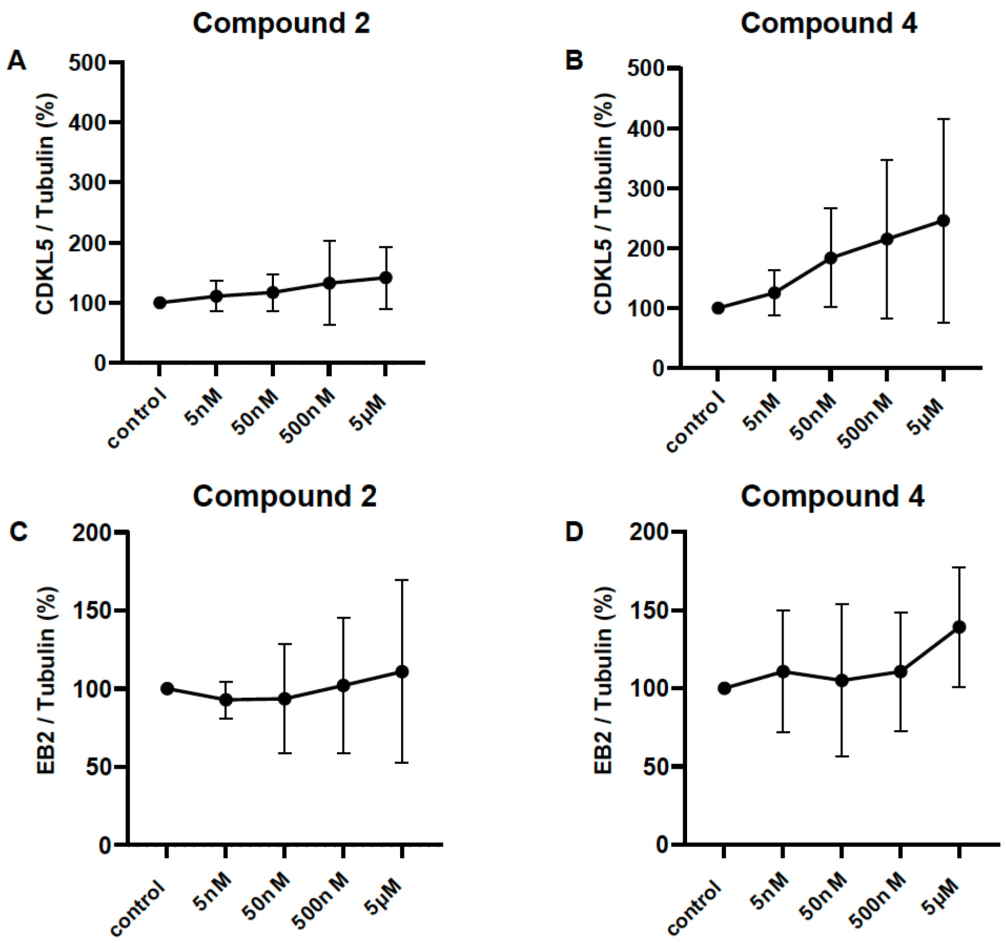


**Figure S5.** CDKL5, EB2 and GSK expression following treatment with compound **2** or **4**. (A, B) Quantification of CDKL5 expression normalized to tubulin following treatment of DIV11-14 rat primary cortical neurons with compound **2** (A) or **4** (B). Two-way ANOVA with pairwise comparisons of each condition against the control condition, non-significant ($p > 0.05$) comparisons are not illustrated. n = 3 replicates. (C, D) Quantification of EB2 expression normalized to tubulin following treatment of DIV11-14 rat primary cortical neurons with compound **2** (C) or **4** (D). Two-way ANOVA with pairwise comparisons of each condition against the control condition, non-significant ($p > 0.05$) comparisons are not illustrated. n = 3 replicates.

**Table S1.** Statistics for CDKL5 data collection, phasing and refinement.

| **Data Collection Statistics** | |
|---|---|
| Radiation source | Diamond I04 |
| Wavelength (Å) | 0.9763 |
| Spacegroup | $P6_322$ |
| Cell dimensions: | |
| $a$, $b$, $c$ (Å) | 88.66 88.66 131.51 |
| $\alpha$, $\beta$, $\gamma$ (°) | 90 90 120 |
| Number of molecules/asymmetric unit | 1 |
| Resolution range (Å) | 44.33-2.201 (2.20-2.24) |
| Total observations | 627548 (29956) |
| Unique reflections | 16108 (759) |
| Completeness (%) | 99.9 (97.9) |
| Multiplicity | 39.0 (39.5) |
| $R_{merge}^{a}$ | 0.181(1.433) |
| Average I/σ *(I)* | 15.5(1.9) |
| $CC_{1/2}$ (%) | 98.6 (92.6) |
| **Refinement and model statistics** | |
| Resolution range (Å) | 44.330 - 2.201 (2.280 - 2.201) |
| Number of reflections used | 16071 (1538) |
| $R_{work}^{b}/R_{free}^{c}$ (%) | 30.98/28.76 |
| Ligand ID | 354 |
| **B values ($Å^2$)** | |
| Overall | 58.200 |
| Macromolecules | 58.700 |
| Ligands | 37.400 |
| Waters | 45 |
| **Root mean square deviation from ideality** | |
| Bond lenghts (Å) | 0.005 |
| Bond angles (°) | 0.904 |
| **Number of atoms** | |
| Protein atoms | 2260.000 |
| Ligands | 30 |
| Waters | 37.000 |
| **Ramachandran analysis** | |
| Favoured / Allowed / Outliers (% of residues) | 94.89/4.74/0.36 |
| Rotamer outliers (%) | 0.430 |
| Clashscore | 12.130 |
| **PDB Code** | 8CIE |

*Values in parentheses are for the highest resolution shell.

**Compound 2**

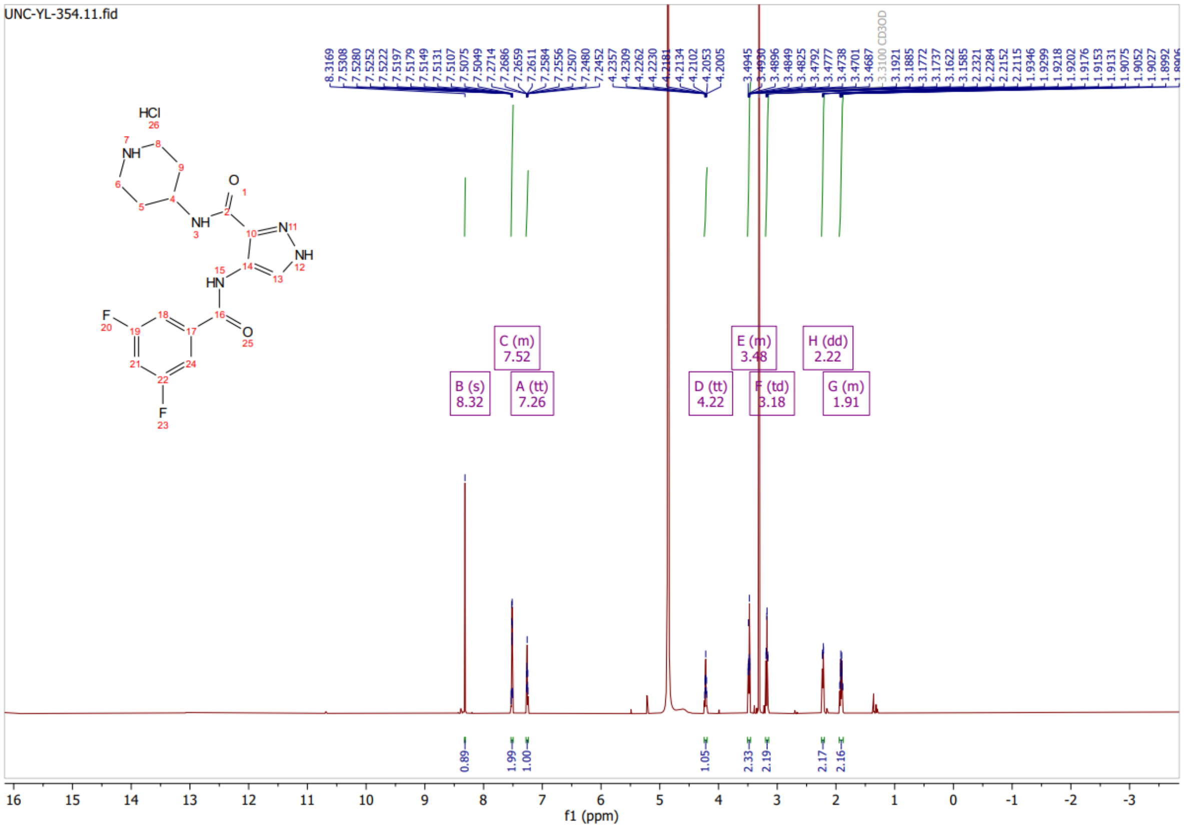

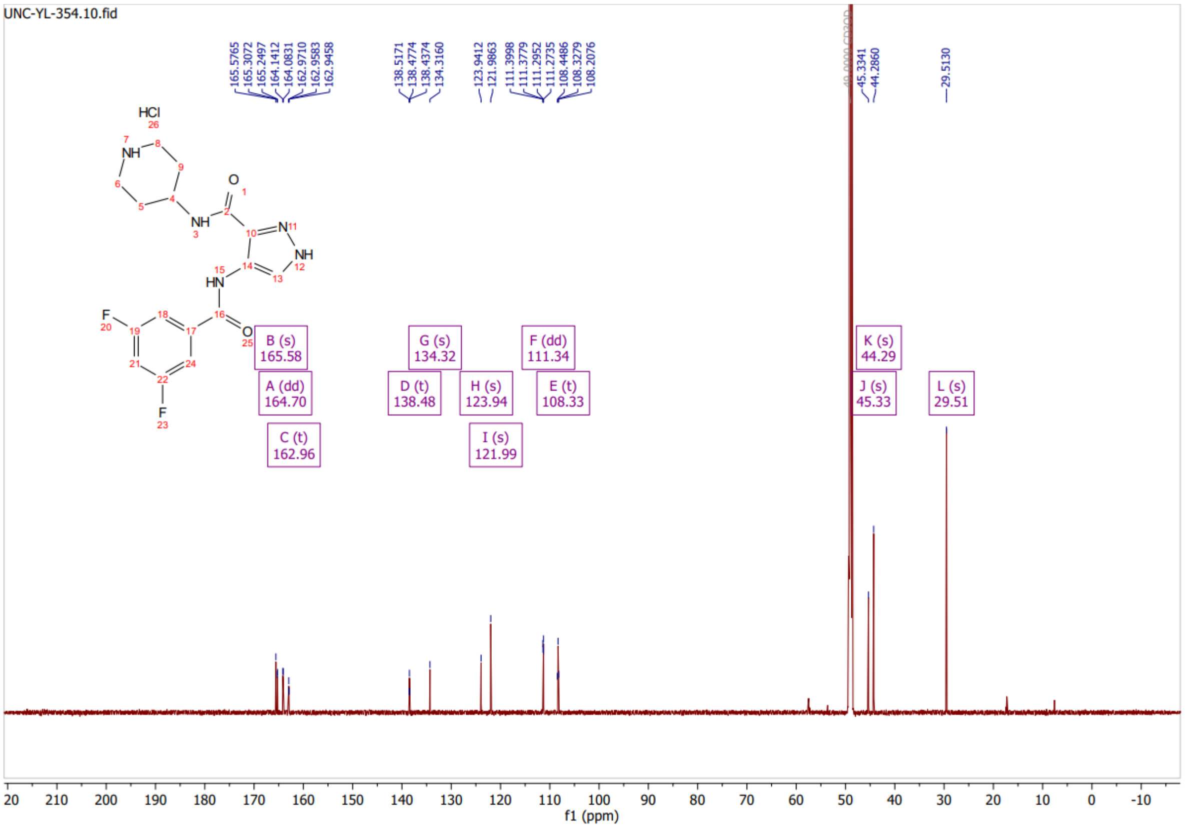

#### Compound 4

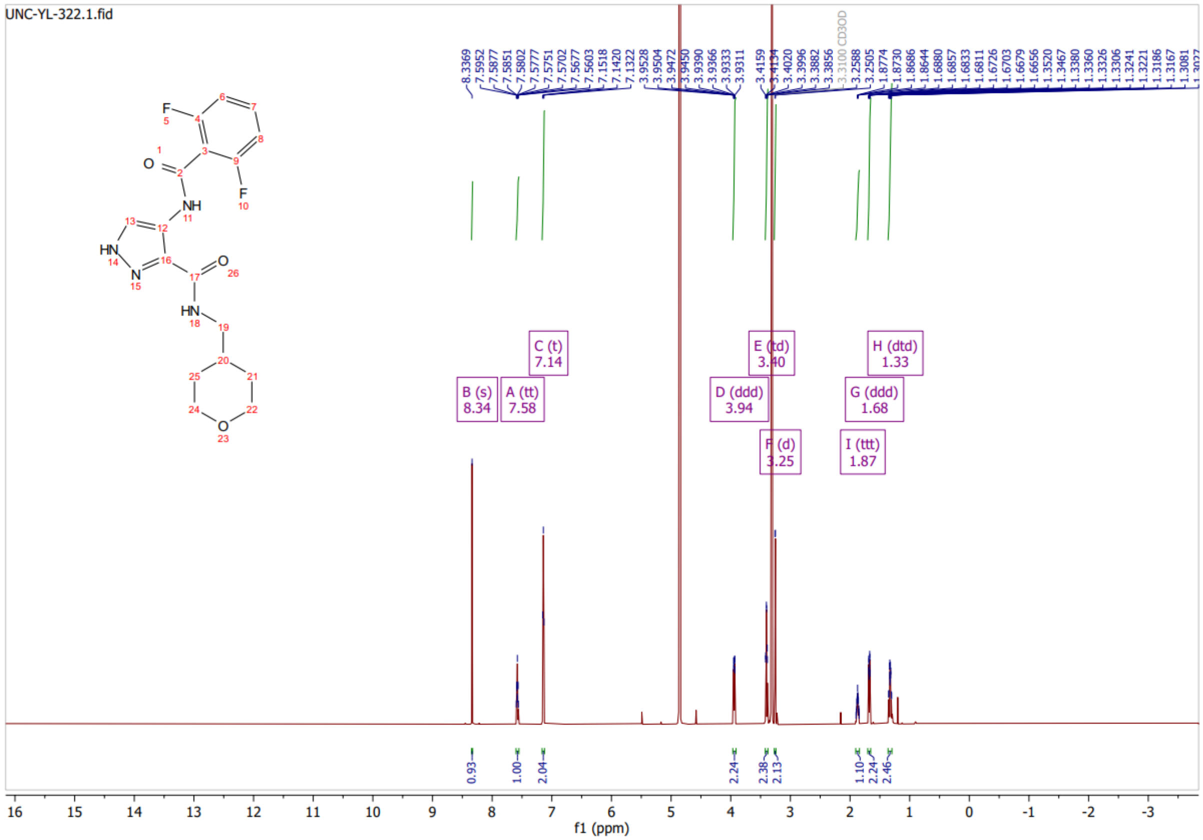

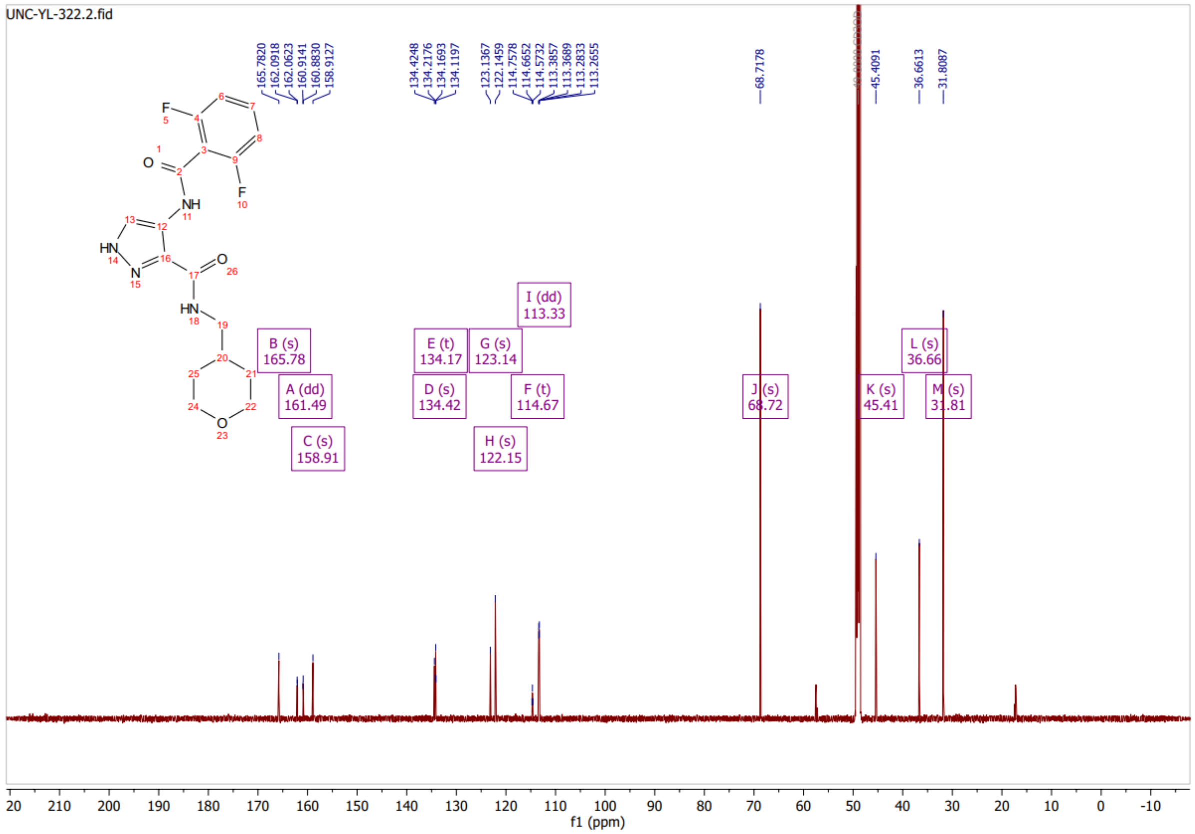
